# Recent trends in Tricolored Blackbird colony size: analysis of the 2008 through 2017 Triennial Statewide Surveys

**DOI:** 10.1101/466532

**Authors:** Timothy D. Meehan, Samantha Arthur, Nicole L. Michel, Chad B. Wilsey, Gary M. Langham

## Abstract

Tricolored blackbird (*Agelaius tricolor*) is a colonial breeder, largely restricted to grasslands, wetlands, and agricultural habitats of California. Tricolored blackbird abundance declined considerably during the 20^th^ century. Recent trends have been less clear, however, hindering efforts to evaluate the conservation needs of the species. We assessed trends in tricolored blackbird colony size using the 2008, 2011, 2014, and 2017 Triennial Tricolored Blackbird Statewide Survey, a community-science effort involving hundreds of volunteer observers. After accounting for variation in observer characteristics and survey effort, we found a clear, statistically significant decline in colony size of approximately –5% per year, which translated to a decrease in colony size of approximately –40% between 2008 and 2017. To the extent that colony size is proportional to total abundance, this decline translates to a species half-life of roughly 15 years. We conclude that tricolored blackbird is in considerable need of protection and recovery efforts.

## INTRODUCTION

Tricolored blackbird (*Agelaius tricolor*) is a colonial nesting species, restricted to wetland, grassland, and agricultural habitats, and a near endemic of California, USA (Beedy et al. 2018). As a colonial nester, tricolored blackbirds are particularly vulnerable to population declines and extinction, as relatively few large colonies comprise a significant proportion of the total population (Cook and Toft 2005). The global abundance of tricolored blackbirds declined considerably over the 20th century, with total counts declining by 89% (Beedy et al. 1991) and average colony size declining by 63% (Graves et al. 2013) from the 1930s through the 1970s. Historically, tricolored blackbirds nested in the large wetland complexes of cattails (*Typha* spp.) and bulrushes (*Schoenoplectus* spp.) that occurred throughout California’s Central Valley (Cook and Toft 2005, Graves et al. 2013, Meese 2014). However, an estimated 95% of wetlands were lost between the 1930s and 1980s (Frayer et al. 1989). Habitat loss and declines in insect prey availability, both consequences of agricultural intensification (Habel and Schmitt 2018), are likely causes of population declines (Beedy et al. 2018).

The restricted geographic range, colonial nesting habit, long-term population decline, and reliance on shrinking wetland habitats makes the tricolored blackbird a species of conservation concern. The species was recently assigned Threatened status under the California Endangered Species Act and is currently being considered for listing under the US Endangered Species Act (Beedy et al. 2018). Status reviews have sought to evaluate whether declines in abundance or colony size have leveled off in recent decades. Graves et al. (2013) found that trends in average colony size between 1980 and 2009 were not statistically significant, possibly due to the heterogeneous nature of the data assembled for analysis. They pointed to the need for more years of standardized data collection before recent trends would become clear.

The Triennial Tricolored Blackbird Statewide Survey (hereafter Triennial Survey) was initiated during the 1980s to collect periodic data on breeding tricolored blackbirds. The Triennial Survey was designed as a community science (a more inclusive term for citizen science) survey that benefits from engagement of skilled volunteers. Since 1994, the survey has been the primary source of information on tricolored blackbird distribution and abundance during the breeding season (Meese 2017). Over the years, the nature of the survey has evolved, with participation increasing and protocols becoming more standardized. Surveys since 2008 are characterized by hundreds of volunteers counting hundreds of thousands of breeding tricolored blackbirds at hundreds of colony sites using a consistent protocol (Kelsey 2008, Meese 2017). Despite the increased effort, standardization, and quality of the survey, inference about the population status of tricolored blackbirds remains challenging. These challenges are similar to those encountered when working with data from other community science monitoring programs, such as US Geological Survey’s North American Breeding Bird Survey (Sauer and Link 2011) and the National Audubon Society’s Christmas Bird Count (Link et al. 2006, Soykan et al. 2016).

Challenges surrounding analysis of data from programs such as the Triennial Survey arise from several sources. First, volunteers, with varying expertise and effort (participation) within and among years, conduct the surveys. When expertise and effort vary randomly, noise is added to data, but when these factors vary systematically, they can bias analysis results (Sauer et al. 1994, Link and Sauer 1999). Second, tricolored blackbird breeding biology makes monitoring their populations a challenge. For example, the species breeds in colonies with upwards of 20,000 individuals, which makes it difficult to estimate precise counts in the field, and produces skewed count distributions that make statistical analysis challenging. Also, colonies are known to move to new locations periodically, so a given sampling location could host 5,000 individuals during one Triennial Survey and none during the next. Tricolored blackbirds also have two breeding pulses, the first in early spring in the southern part of the species range, and a later pulse in the northern part, which necessitates a limited survey window in order to avoid double-counting of birds (Hamilton 1998). These factors present analytical problems that can be remedied, in part, by careful consideration of survey effort, thoughtful data selection, and appropriate choice of statistical methods (Sauer and Link 2011, Hochachka et al. 2012).

Previous research has pointed to several practices that facilitate extracting robust trend estimates from community science data. An established way of dealing with the heterogeneity in observer expertise is to model counts with a statistical model that incorporates random effects for unique observers (Sauer et al. 1994, Sauer and Link 2011). Heterogeneity in observer effort can be accounted for directly by including effort metrics as explicit predictors of counts (Link and Sauer 1999, Soykan et al. 2016). Skewed count distributions can be dealt with effectively by including observation-level random effects (Harrison 2014, 2015).

Here, we report trends in colony size derived from an analysis of count data collected at breeding colonies during 4 Triennial Tricolored Blackbird Statewide Surveys conducted between 2008 and 2017. The 2 goals of the study were to quantify patterns in survey effort and then to model the temporal trend in tricolored blackbird colony size over the study period using established statistical approaches designed to account for variation in survey effort.

## STUDY AREA

Data used for this study came from Triennial Surveys conducted over 44 counties across California, USA (Fig. 1A) (Kelsey 2008, Kyle and Kelsey 2011, Meese 2014, 2017). The majority of birds occurred in colonies located in the San Joachin and Sacramento Valleys, followed by the Central Sierra Foothills and Central Coast regions of the state (Fig. 1A). Colonies were generally located in open semi-natural and agricultural habitats, where common nesting substrates included bulrush (*Schoenoplectus* spp.), cattail (*Typha* spp.), Himalayan blackberry (*Rubus armeniacus*), mallow (*Malva* spp.), mustard (*Brassica* spp.), stinging nettle (*Urtica dioica*) and triticale (*Triticum* × *Secale*) (Kelsey 2008, Kyle and Kelsey 2011, Meese 2014, 2017).

**Figure 1.**
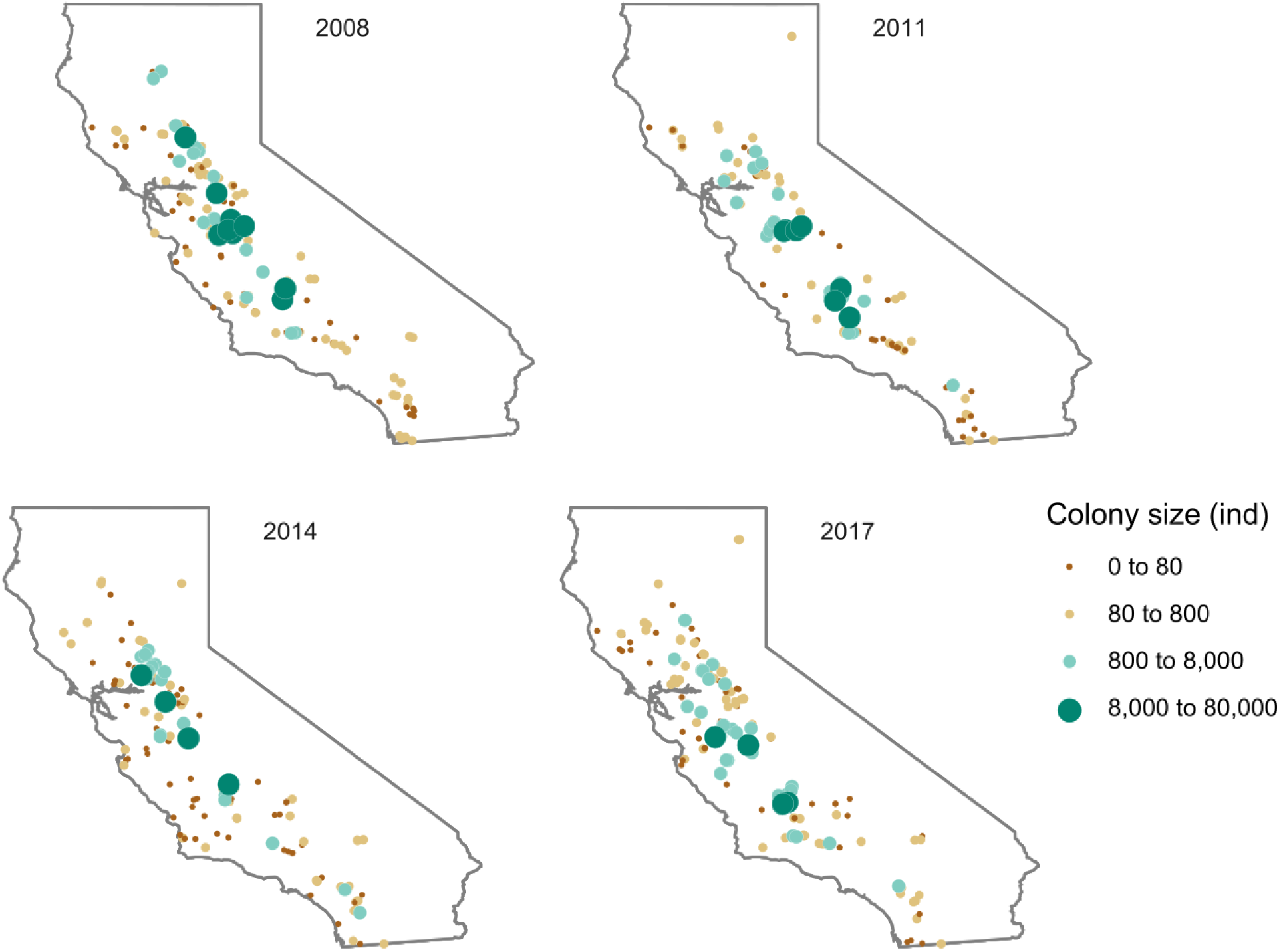
Tricolored blackbird colony sizes (number of individuals) recorded during 4 Triennial Surveys conducted during 2008, 2011, 2014, and 2017 across much of the species geographic range in California, USA.

## METHODS

Triennial Surveys included in this study occurred between 2008 and 2017, a 10-year period in which survey protocols have remained consistent (Kelsey 2008, Kyle and Kelsey 2011, Meese 2014, 2017). Each year, official counts were conducted during a prescribed 3-day window during April. Counts were typically preceded by scouting trips that helped volunteers become familiar with previous colonies and discover new ones. Volunteers counted birds from distances (20 to 100 m) that prevented colony disturbance, and were encouraged to spend approximately 15 min at sites with suitable nesting habitat to determine occupancy. Less time was spent at sites when habitat was obviously unsuitable, and more time was spent as necessary to estimate the number of birds. Full details of the Triennial Survey protocol can be found in annual reports (Kelsey 2008, Kyle and Kelsey 2011, Meese 2014, 2017).

Raw data were obtained for this analysis via the Tricolored Blackbird Portal, a publicly accessible website that serves as a central repository for information on the species (http://tricolor.ice.ucdavis.edu). Before the analysis, we filtered the raw data, keeping only the maximum count per location per year. After filtering, we computed both averaged and summed metrics related to survey effort and blackbird occurrence, to look for systematic patterns of variation among the 4 surveys.

Averaged effort metrics included the mean number of observers participating in the maximum count, per location and year, and the mean duration (min) of the maximum count, per location and year. Summed metrics included the total number of unique observers per year, and the total number of unique locations visited per year. We further decomposed the total number of locations visited into those visited for the first, second, third, and fourth time during the 4 surveys. We also looked for systematic variation in occurrence across the 4 surveys, and explored whether locations were occupied for the first, second, third, or fourth time during the 4 surveys, necessarily ignoring visits and occupancies before 2008.

After assessing patterns in survey effort and occurrence, we assessed temporal trends in the size of tricolored blackbird colonies using a hierarchical modeling approach that is commonly employed for the analysis of count data produced by community science programs (Kéry and Schaub 2011, Sauer and Link 2011, Soykan et al. 2016). For this analysis, we filtered the dataset so that it included all records with a tricolored blackbird count greater than 0. If a location was visited during 2 Triennial Surveys, and it was occupied both times, then both records were included in the analysis. If a location was visited during 2 surveys and it was occupied 1 time, then only the record with a count greater than 0 was included in the analysis. Thus, the dataset included observations from locations occupied only once (singletons) and observations from locations that were occupied repeatedly (repeats). By including all available non-zero counts, the analysis incorporated the greatest information content. The lack of statistical independence among repeats was dealt with in the statistical model used to analyze the dataset, described next.

We modeled tricolored blackbird counts using a hierarchical statistical model that included a temporal trend as well as several terms to account for spatial and temporal variation in survey characteristics (Kéry and Schaub 2011, Sauer and Link 2011, Soykan et al. 2016). The model took the form:

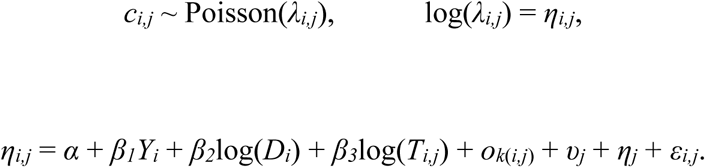

Here, a blackbird count *c* during year *i* at location *j* was drawn from a Poisson distribution with an expected value *λ*_*i*,*j*_. The linear predictor *η*_*i*,*j*_ for the natural log of *λ*_*i*,*j*_ included several fixed effects, including a global intercept *α*, a global log-linear effect *β*_1_ of year *Y_i_*, a global power-law effect *β*_2_ for the number of occupied locations surveyed during a year *D_i_*, and a global power-law effect *β*_3_ for the amount of time spent during the count per location and year *T_ij_*. A log-linear year effect is common in this type of analysis, as it is simple and fits expectations based on population growth (Kéry and Schaub 2011). Flexible power-law effects for the number of occupied locations and survey duration were chosen because it is common for effort effects to be non-linear, with diminishing returns for increased effort (Link and Sauer 1999).

Before analysis, the variables related to global fixed effects were transformed to make units of the global intercept somewhat interpretable. Year was scaled such that 2008 through 2017 was set to 0 through 9. The number of occupied locations surveyed and the duration of surveys were scaled such that minimum values were set to 1. With these transformations, the global intercept represented the natural log of the expected count during 2008, assuming 133 occupied colonies were surveyed for 1 min by a typical observer. Due to the arbitrary nature of these units, expected counts from this analysis should be considered as relative colony size indices (Kéry 2010, Kéry and Schaub 2011, Sauer and Link 2011, Soykan et al. 2016). These indices should not be interpreted as true colony size, although they are thought to be proportional to one another (Link and Sauer 1998, Sauer and Link 2011). However, when confounding factors are properly accounted for, temporal trends in indices are expected to follow trends in true colony size (Link and Sauer 1998, Sauer and Link 2011).

The linear predictor also included four exchangeable random effects, including an exchangeable random intercept per the *k*^th^ observer *O*_*k*(*i*,*j*)_, an exchangeable random intercept per location *ν_j_*, an exchangeable random log-linear effect of year per location *η_j_*, and an overdispersion term *ε*_*i*,*j*_. Random effects were modeled with normal distributions with means = 0 and precisions (inverse of variances) *τ_o_*, *τ_ν_*, *τ_η_*, and *τ_ε_*, respectively. Note that the random observer effect was added to account for variation in counts related to volunteer experience and ability (Sauer et al. 1994, Sauer and Link 2011). Random intercepts and year effects across locations were added to allow the year effect to vary across locations and to account for the correlation between repeat measurements (Zuur et al. 2009). The overdispersion term was added so that overdispersed count data could be modeled using the Poisson distribution (Harrison 2014).

Inference on model parameters followed a Bayesian framework as implemented in the R-INLA package (Rue et al. 2017) for R statistical-computing software (R Version 3.3.1, www.rproject.org, accessed 21 June 2016). Prior distributions for global fixed effects were normal distributions with mean = 0 and precision = 0.001. Priors for *τ_o_*, *τ_ν_*, *τ_η_*, and *τ_ε_* were gamma distributions with parameter values shape = 1 and rate = 0.5. Inference was drawn using 20,000 samples from the joint posterior distribution. In some cases, full marginal posterior distributions were shown in figures. When numerical summaries were given for estimates of model parameters, point estimates were posterior medians with symmetric 95% credible intervals bounded by the 0.025^th^ and 0.975^th^ quantiles of marginal posterior distributions.

## RESULTS

The raw data consisted of 2,713 records from 1,455 locations across 4 annual surveys spanning 10 years. Annual means for the number of observers participating in the maximum count per location, and the duration of the maximum count per location, did not appear to vary systematically across the 4 surveys (Table 1). In contrast, the total number of observers conducting surveys, and the total number of locations visited, increased by approximately 25% and 150%, respectively (Table 1). Locations visited during the 2011 through 2017 surveys were a combination of new locations and previously visited ones (Table 1). Across the 4 surveys, between 296 and 416 locations were visited for the first time, between 196 and 289 were visited for a second time, and between 131 and 219 were visited for a third time. During 2017, 103 locations were visited for a fourth time (Table 1).

**Table 1.**
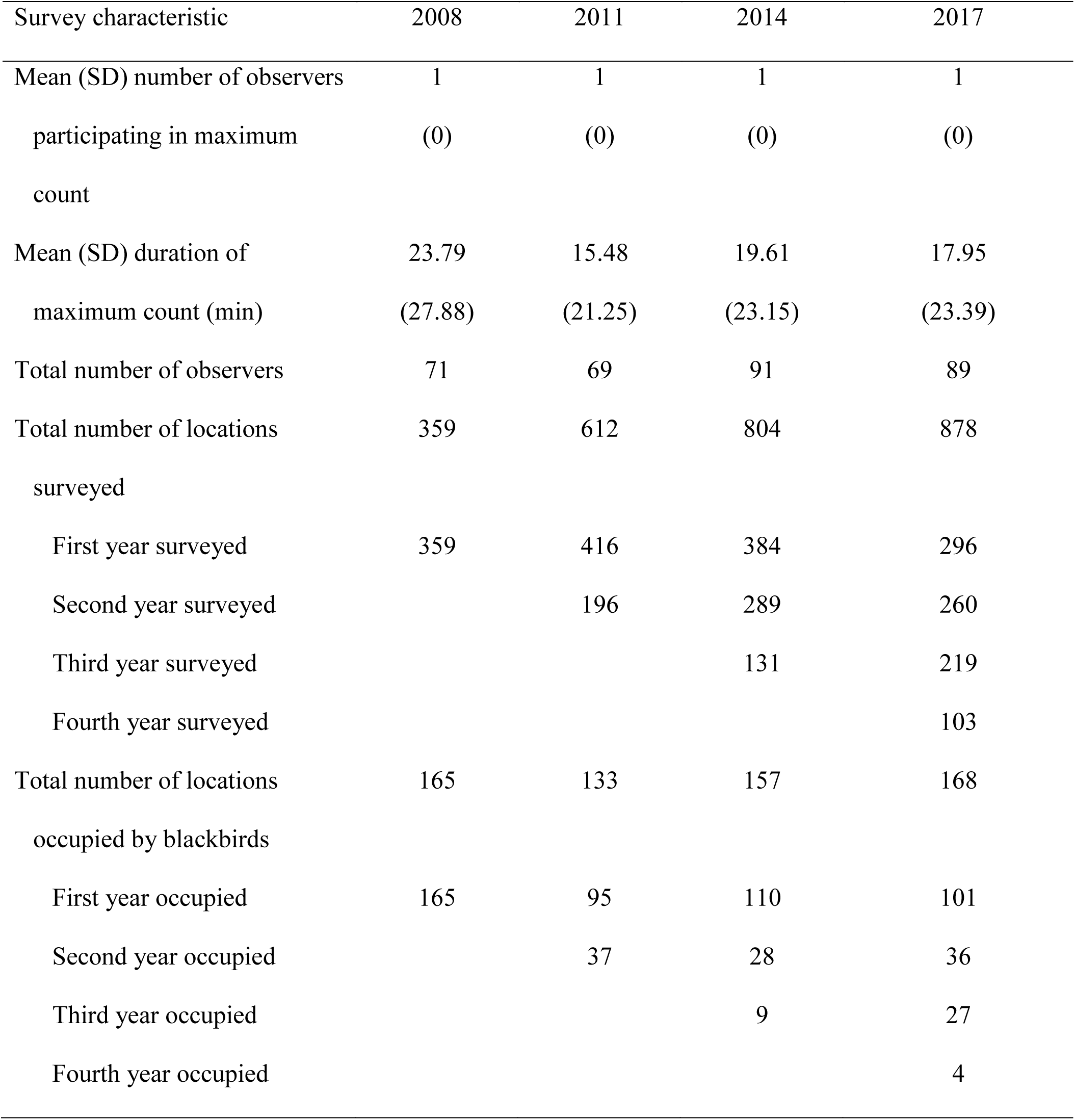
Summary of effort and occurrence during 2008, 2011, 2014, and 2017 Tricolored Blackbird Triennial Surveys.

Despite the increasing numbers of total observers and number of locations visited, there was not systematic variation in the number of locations with tricolored blackbirds detected (Table 1). Indeed, blackbirds were detected at 165 locations in 2008 and 168 locations during 2017. Across the 4 surveys, between 95 and 165 locations were occupied for the first time, between 28 and 37 were occupied for a second time, and between 9 and 27 were occupied for a third time. During 2017, 4 locations were occupied for a fourth time (Table 1).

Of the 1,455 locations visited during the study period, 710 were visited during 1 year, while 395 were visited during 2 years, 247 were visited during 4 years, and 103 were visited during 4 years of the survey. Of the 1,198 possible state transitions in the dataset (i.e., 395 + [247 × 2] + [103 × 3]), there were 746 cases (62%) where a location remained unoccupied, 90 cases (8%) where a location transitioned from unoccupied to occupied, 256 cases (21%) where a location transitioned from occupied to unoccupied, and 106 cases (9%) where a location remained occupied. Of the 452 transitions (90 + 256 + 106) where at least 1 state was occupied, 20% of transitions were from unoccupied to occupied, 57% of transitions were from occupied to unoccupied, and 23% of transitions were from occupied to occupied.

Of the 1,455 locations visited during the study period, 471 had blackbirds detected at least 1 time. The 612 counts from these 471 locations suggested that colony size had declined significantly over the 4 surveys. This trend is apparent, qualitatively, in Figure 1, as the number of large green points indicating large colonies declined between 2008 and 2017, and statistically, in Table 2, from the count model analysis.

**Table 2.** Summaries of marginal posterior distributions for global fixed effects from the hierarchical model of colony size counts.

|  | Posterior | Lower | Upper | Probability |
| --- | --- | --- | --- | --- |
| Model Parameter | median | credible limit | credible limit | estimate $\geq 0$ |
| Global intercept, $\alpha$ | 3.951 | 3.339 | 4.554 | $> 0.999$ |
| Survey year, $\beta_1$ | -0.050 | -0.106 | 0.006 | 0.038 |
| $\log(\text{Locations occupied}), \beta_2$ | -0.052 | -0.178 | 0.075 | 0.212 |
| $\log(\text{Count duration}), \beta_3$ | 0.437 | 0.285 | 0.590 | $> 0.999$ |

Output from the model analysis gave a posterior median of −0.050 for *β*_1_ (the effect of survey year on expected counts), with a 95% credible interval of −0.106 to 0.006 (Table 2). While the symmetric credible interval included 0, the posterior probability that *β*_1_ was ≥ 0 was 0.038. The posterior median converted to a percent population decline of approximately −5% per year (Figure 2). When scaled over the 10-year study period, that annual decline translated to an approximate 40% decrease in colony size between 2008 and 2017.

**Figure 2.**
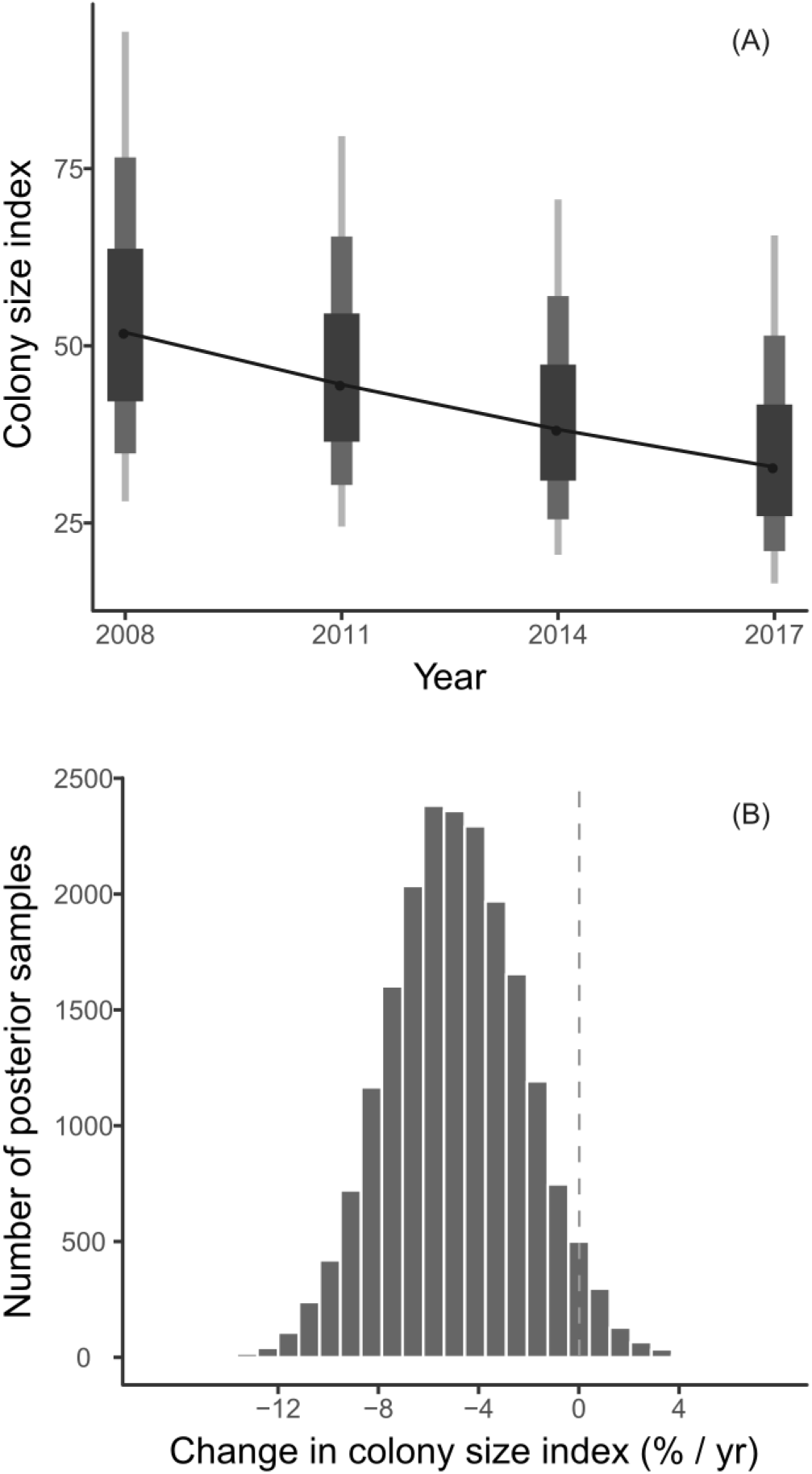
Posterior probabilities for estimates of (A) colony size index and (B) annual percent change in colony size index, derived from analysis of counts conducted at tricolored blackbird colonies during 2008, 2011, 2014, and 2017 across much of the species geographic range in California, USA. In the left panel, vertical lines, from widest to narrowest, represent the middle 50%, 80%, and 95% of posterior distributions, respectively.

The posterior median value for *β*_2_ (the effect of the number of occupied locations) was −0.052 (−0.178, 0.075; Table 2). Given the wide credible interval encompassing 0, there was little evidence that observed blackbird counts were related to the number of occupied colonies surveyed in a given year. The posterior median value for *β*_3_(the effect of survey duration) was 0.437 (0.285, 0.590; Table 2). Given that the credible interval did not encompass 0, there was evidence for a positive relationship between observed blackbird counts and count duration. The value of 0.437 could be interpreted as an exponent for an effort-correction function with a power law form. The exponent between 0 and 1 indicated a positive effect of count duration, but that continued increases in count duration yielded diminishing returns.

Regarding random effects, median values for *τ_o_*, *τ_ν_*, *τ_η_*, and *τ_ε_* were 1.082, 2.226, 27.877, and 0.431, respectively. These values, converted to a standard deviation scale, were 0.961, 0.670, 0.189, and 1.523, respectively. Credible intervals for all forms of *τ* did not include 0, so all random effects were useful components of the model. In terms of highest to lowest variation explained, random effects ranked *ε*, *o*, *ν*, and *η* Thus, the random observer effect *o* explained more variation than the location effect *u* and the random year effect per location *η*. The importance of *ε* indicated that the colony counts were indeed overdispersed relative to a Poisson distribution.

Posterior distributions for the model intercept and the global year effect (Table 2) were used to compute an annual tricolored blackbird colony size index, *N_i_* = exp(*α* + *β*_1_*Y_i_*). Figure 2B illustrates the magnitude of the model-estimated decrease in the colony size index from 2008 through 2017.

## DISCUSSION

It is clear that tricolored blackbird populations in California declined considerably during the 20^th^ century (Beedy et al. 1991, Graves et al. 2013). Analysis of more recent trends, using heterogeneous data collected from the peer-reviewed and gray literature, has yielded results that are less clear (Graves et al. 2013), hindering efforts to evaluate the conservation needs of the species. We explored recent trends in colony counts conducted by community volunteers, using an established analytical approach that takes into account variation in observer characteristics and survey effort. After accounting for these potentially confounding variables, there was a clear, statistically significant decline in colony size of approximately 5% per year, which translated to a decrease in colony size of approximately 40% between 2008 and 2017.

The tricolored blackbird is listed as Endangered by the International Union for Conservation of Nature. The species was recently assigned Threatened status under the California Endangered Species Act, and is currently being considered for listing under the US Endangered Species Act (Beedy et al. 2018). The Partners in Flight Landbird Conservation Plan lists the species on its Red Watch List, with an estimated species half-life of 50 years or more (Rosenberg et al. 2016). To the extent that colony size is proportional to total abundance, the 5% annual decline, documented in this analysis, suggests a species half-life closer to 15 years than 50 years. Thus, the outlook for this species may be worse than assumed.

We quantified observer effort over the 10-year study period and found that some metrics varied considerably (count duration) while others varied little (number of observers per count), and that some varied systematically (total number of observers and locations visited) while others did not (total number of active colonies counted). This variation could have implications for observed trends if not properly accounted for (Sauer et al. 1994, Link and Sauer 1999). For example, over the study period, the number of volunteer observers increased substantially. A systematic change in average observer ability, as new, inexperienced observers joined the survey, could have resulted in an apparent trend in blackbird counts. We reduced this possibility by including observer identity directly into the model. Second, the number of sites visited increased substantially over the study period. Despite this increase, the number of locations with blackbird detections remained remarkably consistent. It was conceivable that an apparent decline in colony size could be driven by a gradual increase in the number of colonies counted, under the assumption that large colonies would be found and counted before small ones. This possibility was not likely to occur in this study, as there was not systematic variation in the number of counted colonies across the study period, and because the number of occupied colonies was incorporated directly into the model. Third, the amount of time spent during colony counts varied considerably within and across surveys. Systematic declines in count duration across years could have resulted in an apparent decrease in colony size. This possibility was unlikely to influence our results, as count duration was explicitly modeled in our analysis.

## MANAGEMENT IMPLICATIONS

The main management implications from this study are that the tricolored blackbird continues to decline in California, and that current conservation status and management activities have been insufficient to stop this decline. Our study does not address how this decline might be stopped. However, the 5% annual decrease in colony size observed during this study was statistically indistinguishable from the 6% decline in overall abundance recently reported from a study (Robinson et al. 2018) that analyzed completely different count data from the eBird program (Sullivan et al. 2014). The similarity in annual declines across the two studies suggest that tricolored blackbird population decline is occurring through a decline in colony size, possibly more so than a decline in colony number. The integrated population model analysis by Robinson et al. (2018) combined eBird relative abundance data with independent data on productivity and survival. Correlations between population growth rates and vital rates suggested that population growth was especially sensitive to adult female survival and productivity (Robinson et al. 2018). They suggested that management activities should focus on increasing annual productivity, as adult female survival is already relatively high. Conservation programs, such as payments to farmers in California’s Central Valley to delay harvest of silage and grain crops when fields are occupied by tricolored blackbird nestlings (Arthur 2015), that increase productivity could reverse the negative trends in colony size observed in the Triennial Survey.

## ACKNOWLEDGMENTS

We acknowledge and thank the hundreds of community volunteers who have participated in the Triennial Tricolored Blackbird Statewide Survey for producing an expansive, high-quality dataset on tricolored blackbird colony size. We thank E. C. Beedy, R. J. Meese and M. Holyoak for sharing the data, and for lending insight into how they were collected.

---

**Figure S1.**
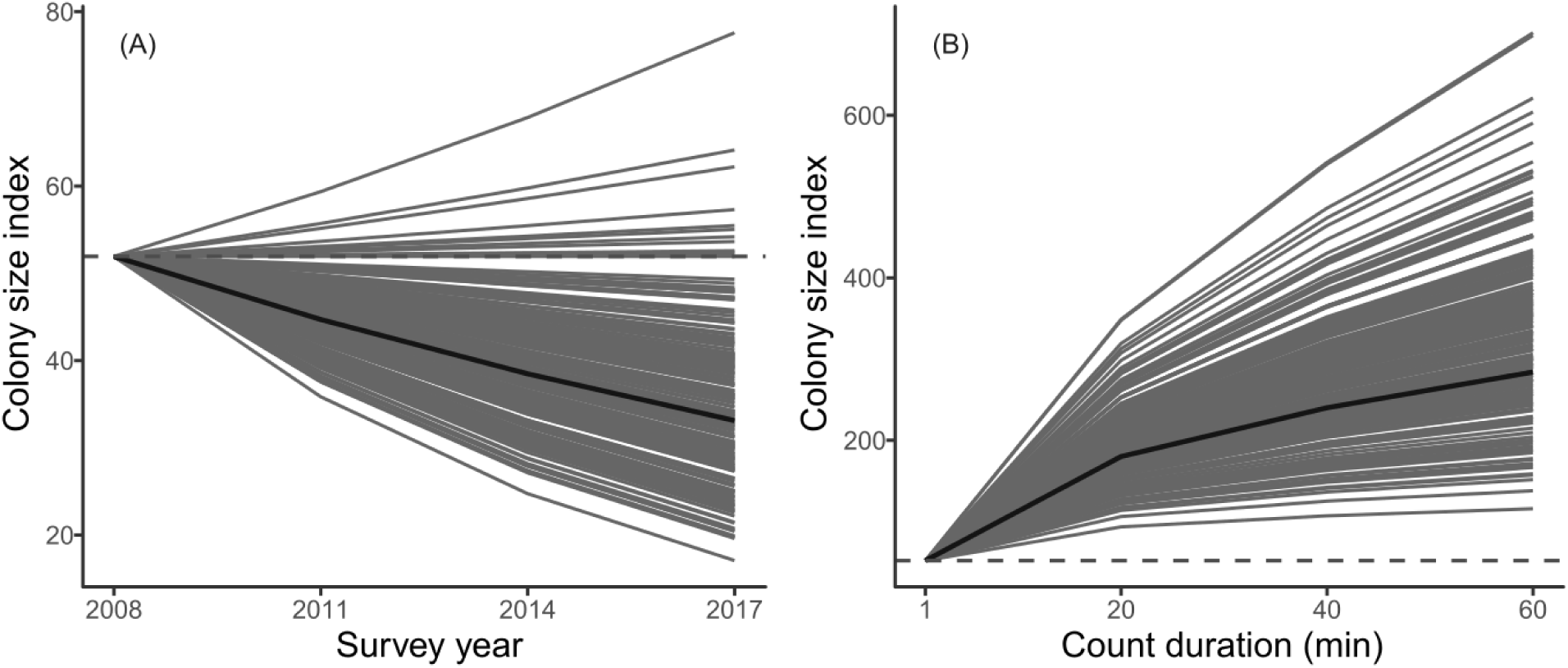
Partial plots for the effects of (A) year (2008 through 2017) and (B) count duration (min) on counts at Tricolored Blackbird colonies. The posterior median effects are shown with dark brown lines, while light brown lines reflect 200 random draws from effect posteriors.

**Figure S2.**
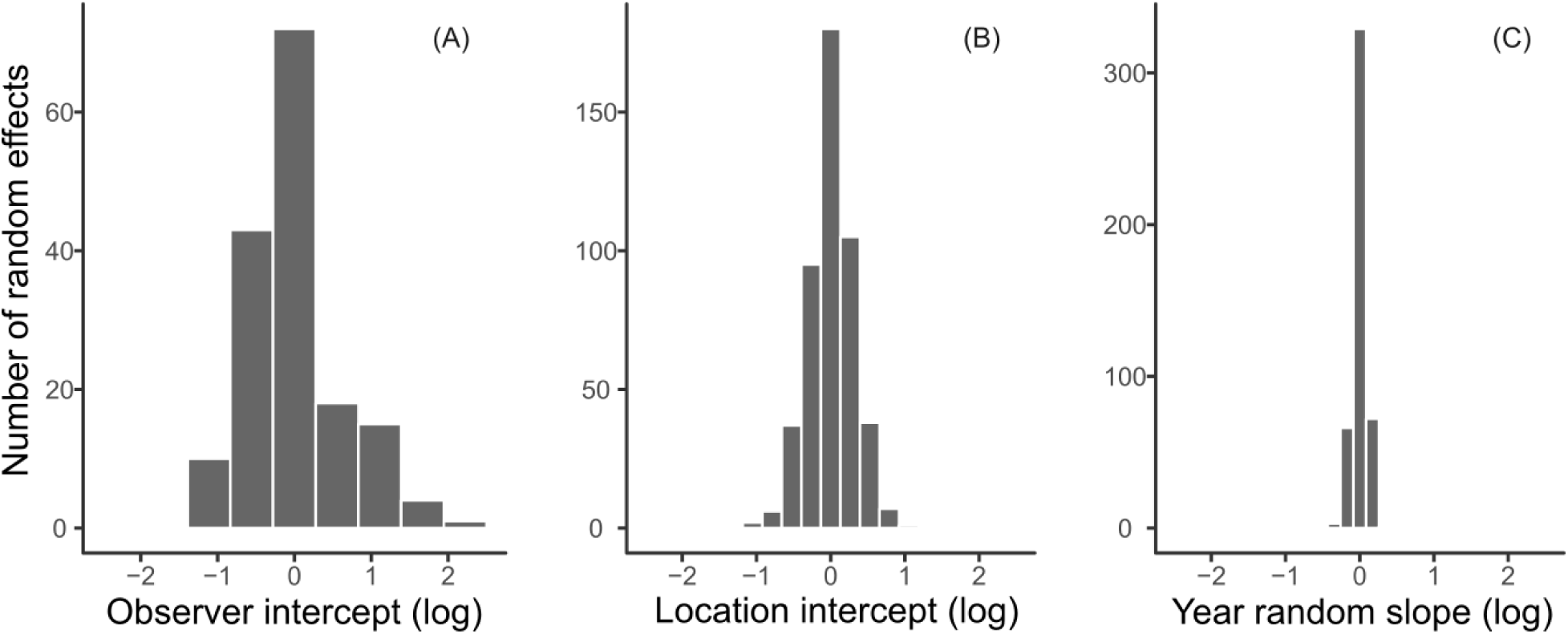
Histograms depicting the variation associated with three random effects incorporated into the count analysis model. The x-axes are scaled identically to facilitate comparison.

